# Transcription-replication conflicts as a source of common fragile site instability caused by BMI1-RNF2 deficiency

**DOI:** 10.1101/846683

**Authors:** Anthony Sanchez, Angelo de Vivo, Peter Tonzi, Jeonghyeon Kim, Tony T. Huang, Younghoon Kee

## Abstract

Common fragile sites (CFSs) are breakage-prone genomic loci, and are considered to be hotspots for genomic rearrangements frequently observed in cancers. Understanding the underlying mechanisms for CFS instability will lead to better insight on cancer etiology. Here we show that Polycomb group proteins BMI1 and RNF2 are suppressors of transcription-replication conflicts (TRCs) and CFS instability. Cells depleted of BMI1 or RNF2 showed slower replication forks and elevated fork stalling. These phenotypes are associated with increase occupancy of RNA Pol II (RNAPII) at CFSs, suggesting that the BMI1-RNF2 complex regulate RNAPII elongation at these fragile regions. Using proximity ligase assays, we showed that depleting BMI1 or RNF2 causes increased associations between RNAPII with EdU-labeled nascent forks and replisomes, suggesting increased TRC incidences. Increased occupancy of a fork protective factor FANCD2 and R-loop resolvase RNH1 at CFSs are observed in RNF2 CRISPR-KO cells, which are consistent with increased transcription-associated replication stress in RNF2-deficient cells. Depleting FANCD2 or FANCI proteins further increased genomic instability and cell death of the RNF2-deficient cells, suggesting that in the absence of RNF2, cells depend on these fork-protective factors for survival. These data suggest that the Polycomb proteins have non-canonical roles in suppressing TRC and preserving genomic integrity.

**Author summary:** Increasing evidence suggest that instabilities at common fragile sites (CFSs), breakage-prone genomic loci, may be source of genomic aberration seen in cancer cells. Among the proposed mechanisms that can cause CFSs instabilities is the conflict between transcription and replication, and the mechanisms or factors that resolve the possible conflicts are only beginning to be understood. Here we found that deficiency in the Polycomb group proteins BMI1 or RNF2 leads to the CFS instability, and is associated with transcription-associated replication fork stresses. We further found that in the absence of RNF2, cells depend on the Fanconi Anemia fork-protective proteins for genome maintenance and survival. These results underscore that the Polycomb proteins are important for genome maintenance.

## Introduction

Common fragile sites (CFSs) are natural genomic loci that are prone to gaps and breaks upon DNA replication stress. CFSs are hotspots for genomic aberrations, which are frequently found in cancerous cells (1). It is generally accepted that perturbations in DNA replication may be the underlying cause for CFS instability. A few potential sources for the replication defects and increased breakages at CFSs have been proposed, such as high frequencies of DNA secondary structures forming barriers to the fork progression, scarcity of replication origins, and collisions between transcription and replication (2–5).

Many CFSs harbor long genes in which transcription and replication can occur simultaneously, elevating the chance for transcription-mediated interference of the replication fork progression (6–9). Prokaryotic and eukaryotic cells appear to have evolved mechanisms to prevent transcription-replication conflicts (TRCs), by separating the timing and location of transcription or replication processes (10). This may be particularly challenging at long genes where the transcription of a single long gene can take place throughout the entire cell cycle including when replication is active (8). TRC incidences can be also accelerated by the overexpression of oncogenes such as MYC (11), RAS (12), or Cyclin E (13), which could alter replication origin firing or global transcription.

TRCs are generally associated with increased levels of R-Loops, a form of RNA-DNA hybrid with a displaced single-stranded DNA, which could aggravate replication fork stalling and DNA breakages (14, 15). Increasing evidence points to TRCs and R-loops as serious threats to genomic stability. Factors that suppress the TRCs and R-loop formation are only beginning to be understood; for one example, recent studies highlighted the role of Fanconi Anemia proteins in recognizing and suppressing R-loops and preserving CFS stability (16–20).

BMI1 and RNF2 are core members of the Polycomb Repressive Complex 1 (PRC1) transcriptional repressors that maintain chromatin in a silenced state. They are required for stem cell maintenance and also have been implicated in cancer development (21). PRC1 induces gene silencing in part by catalyzing histone H2A ubiquitination (H2AK119-ub) or by inducing chromatin compaction (21). Purified RING domains of BMI1 and RNF2 form a heterodimer and induce the H2A ubiquitination, in which the RING domain of RNF2 provides the catalytic activity (E2 binding) and BMI1 serves as a stimulating co-factor (22, 23). In addition to the role in targeted gene silencing and stem cell maintenance, BMI1 and RNF2 also participate in genome stability maintenance; BMI1 localizes to DNA breaks and facilitates DNA repair factors recruitment (24–28). RNF2 also localizes to DNA damage sites where it induces nucleosome remodeling (29). Several studies showed that BMI1 depletion causes uncontrolled transcription at nuclease and UV-induced DNA double strand breaks (DSBs) (30–32), suggesting that one way BMI1 promotes genome stability is by controlling RNAPII elongation at the DNA lesions. These findings suggest that BMI1 and RNF2 can directly promote genome stability independently of targeting specific gene repression, and led us to hypothesize that loss of the RNAPII-controlling activity of BMI1 and RNF2 may cause transcription-induced instability in breakage-prone loci such as CFSs.

Here we show that BMI1 and RNF2 are important for preserving CFS stability. Depletion of BMI1 or RNF2 causes increased replication stress and fork stalling. The replication defects are associated with deregulated RNAPII activities; increased RNAPII occupation is observed in CFSs, and the physical coupling of replisome and the Pol II complex is observed in cells depleted of BMI1 and RNF2, which can be reversed by inhibiting RNAPII elongation. Consistently, BMI1 or RNF2 depleted cells exhibit increased fork stalling and reduced rate of replication at CFSs. We found that CFSs in RNF2 KO cells are more enriched with FANCD2 and RNH1, both of which are required to resolve R-loops. Depleting FANCD2 or FANCI in RNF2 KO cells synergistically increased the genomic instability, further suggesting the important roles of the FA proteins in responding to the R-loop-associated CFS instability. Altogether, our work provides an insight into the role of Polycomb components in suppressing genomic aberration, which is distinct from its canonical role in epigenetic silencing linked to cell stemness maintenance.

## Results

### Depletion of BMI1/RNF2 leads to CFS instability and replication fork stress

We first took notice that cells depleted of BMI1 or RNF2 display retardation in the cell cycle progression, upon release from synchronization at the G1/S boundary by HU (Fig 1A). The retarded progression through S phase may indicate that these cells experience replication stress. Depletion of factors that mitigate replication stress often leads to fragilities or breakages at common fragile sites (CFSs). Studies found that 53BP1-containing nuclear bodies (53BP1-NBs) in G1 cells at CFSs result from replication stress from previous generation (33, 34). We found that siRNA-mediated silencing of BMI1 or RNF2 in U2OS cells leads to consistent and notable increase in the 53BP1-NBs, particularly in cyclin A-negative G1 cells (Fig 1B). We found that the extent of 53BP1-NB formation was comparable to the silencing of FANCD2, a replication fork-associated protein required for normal replication fork elongation, and slightly less than the silencing of topoisomerase TOP2A. The 53BP1-NBs are further increased when the cells were treated with replication stressor Aphidicolin (APH) (representative images are in S1 Fig). The increase by RNF2 depletion was reversed to the control level by re-expressing RNF2 (Fig 1C). To investigate the phenotype further, we introduced a CRISPR Cas9-mediated RNF2 knockout (KO) in untransformed human ovarian epithelial T80 cells (S2 Fig). We noted that the KO cells exhibit slower growth rate compared to parental T80 cells yet maintained the ability to form colonies and survived until ∼10 passages. At early passages, we consistently found that 53BP1-NBs are increased in the RNF2 KO cells (S3 Fig). The occupation of 53BP1 at CFSs was confirmed by anti-53BP1 chromatin immunoprecipitation (ChIP) at several prominent CFS loci (schematic in Fig 1D), in the CRISPR Cas9 RNF2 KO cells (Fig 1E) or in siRNA-mediated knockdown cells (Fig 1F). It was notable that the 53BP1 occupation fold change in the absence of RNF2 was comparable to that of APH-treated T80 cells (S4 Fig). These results also confirm that the tested CFSs are “expressed” in T80 cells, when replication stress is induced. Cells depleted of factors that regulate replication stress have difficulty recovering from HU treatment (35). Consistently, BMI1 or RNF2 knockdown increased cellular sensitivity to HU and APH (Figs 1G and 1H). Incomplete replication at CFSs is known to lead to formation of ultrafine bridges during Anaphase, aberrant mitotic structures, and micronuclei (4, 36). Consistent with this notion, we observed increased presence of micronuclei in both BMI1 and RNF2-depleted cells (Fig 1I).

**Figure 1.**
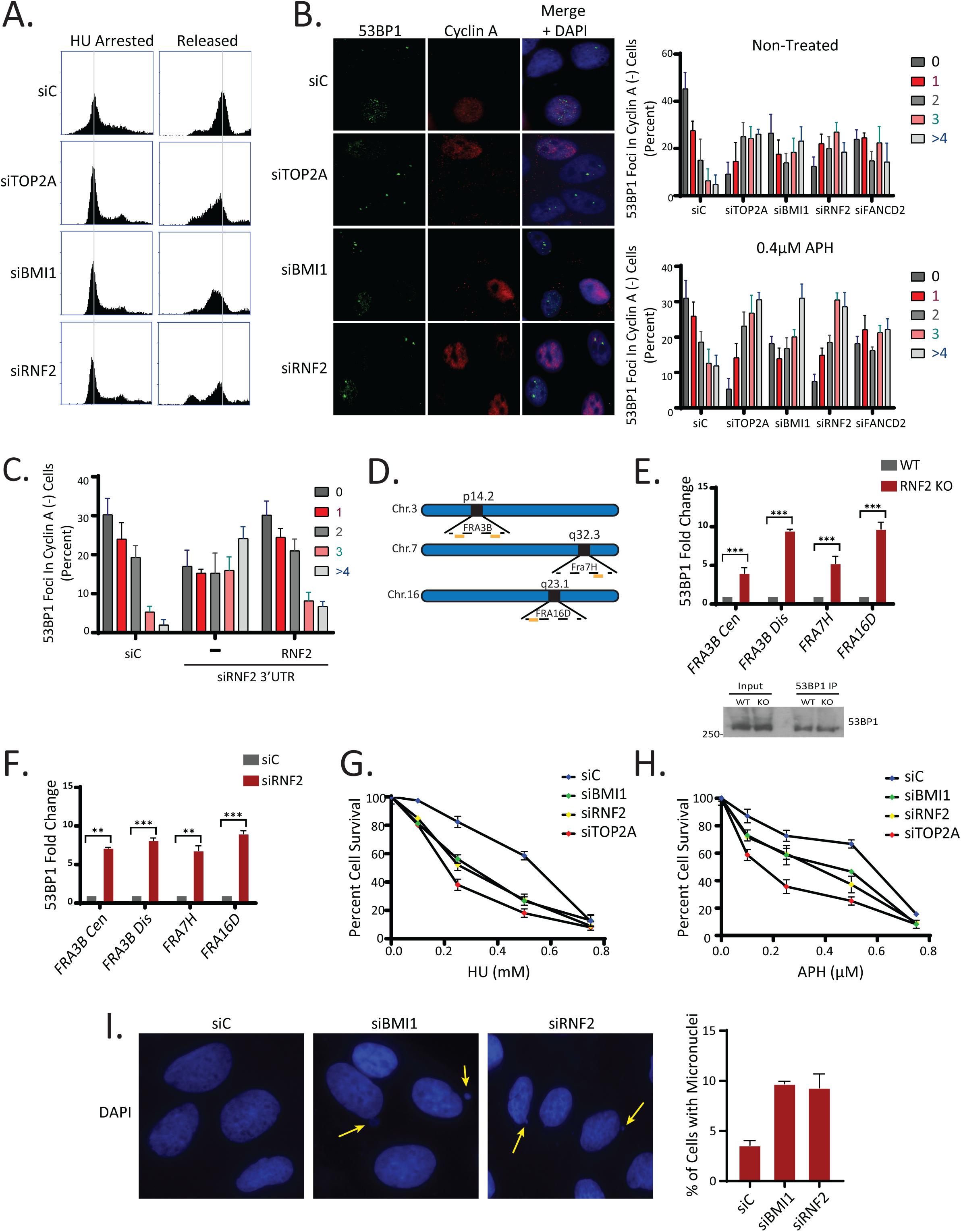
BMI1 and RNF2-deficiency results in replication-associated instabilities. **A.** HU-arrested U2OS cells show delayed cell cycle progression when BMI1 or RNF2 is depleted by siRNA. siTOP2A was used as a positive control (N=3 biological replicates). **B.** (Left) Representative images of U2OS cells stained with 53BP1 and Cyclin A following treatment with siRNAs for control (scrambled), BMI1, RNF2 and TOP2A. (Right) quantification of 53BP1 foci in Cyclin A negative cells (N=100 from 3 biological replicates). **C.** U2OS cells transfected with siRNAs were subsequently transfected with a 3xFLAG-RNF2 plasmid. At 72 hours post-transfection, cells were fixed and analyzed for 53BP1 foci (in Cyclin A-negative cells). Assays were done in triplicates (N=100 for each condition). **D.** Schematic for CFS primer binding locations on FRA3B, FRA7H and FRA16D used in ChIP experiments. **E.** (Top) qPCR quantification of 53BP1 ChIP in T80 wild type and RNF2 KO cells (N=3 biological replicates; ***P <0.0005, **P <0.005). (Bottom) Western blot confirmation of 53BP1 IP in wild type and RNF2 KO cells. **F.** qPCR quantification of anti-53BP1 ChIP in T80 cells transfected with either control or RNF2 siRNAs (N=3 biological replicates; ***P <0.0005, **P <0.005). **G.** Clonogenic survival assay determines that T80 cells depleted of BMI1, RNF2 or TOP2A by siRNAs are sensitive to treatment with HU. (N=3 biological replicates). **H.** Clonogenic survival assay determines that T80 cells depleted of BMI1, RNF2 or TOP2A by siRNA are sensitive to treatment with Aphidicolin (APH) (N=3 biological replicates). **I.** (Left) Representative images showing that U2OS cells depleted of BMI1 or RNF2 by siRNAs harbor increased micronuclei. (Right) Quantification of the percentage of cells with micronuclei. (N=50 from 3 biological replicates).

### Depletion of BMI1 or RNF2 causes replication fork stresses

Since replication stress is a major cause for the CFS instability, we wished to test if BMI1 or RNF2 depletion affects replication fork elongation in unperturbed conditions. To do so, we used the DNA fiber assay to assess DNA replication elongation rate (speed) by labeling the asynchronously growing RPE1 cells that were treated with each siRNA. Cells were initially pulse-labeled with iododeoxyuridine (IdU) to mark all elongating replication forks, followed by a wash step and addition with fresh media containing chlorodeoxyuridine (CldU) (see DNA fiber labeling schematics, Fig 2A). Quantitative analysis of individual CldU-labeled track length revealed statistically significant decrease in DNA replication speed in both BMI1 and RNF2-depleted cells compared to control cells (Fig 2A). These results demonstrate that both BMI1 and RNF2 are important for maintaining the proper elongation rate of DNA replication. We further noted that patterns of bidirectional fork movement are more asymmetric in both BMI1 and RNF2-depleted cells; while control cells showed that approximately 13% of the fibers displayed asymmetry, 43% and 43% of the fibers for two BMI1 siRNAs and 38% and 38% for two RNF2 siRNAs displayed the asymmetry (Fig 2B). The differences in the fork speed between the two forks in each fiber suggest that BMI1 or RNF2 deficiency causes the forks to stall due to some physical obstacles (e.g. transcriptional collisions or damaged DNA), rather than due to some global influences (e.g. overall cell growth change or nucleotide deprivation). Altogether, these results support that BMI1 or RNF2 deficiency causes replication stresses.

**Figure 2.**
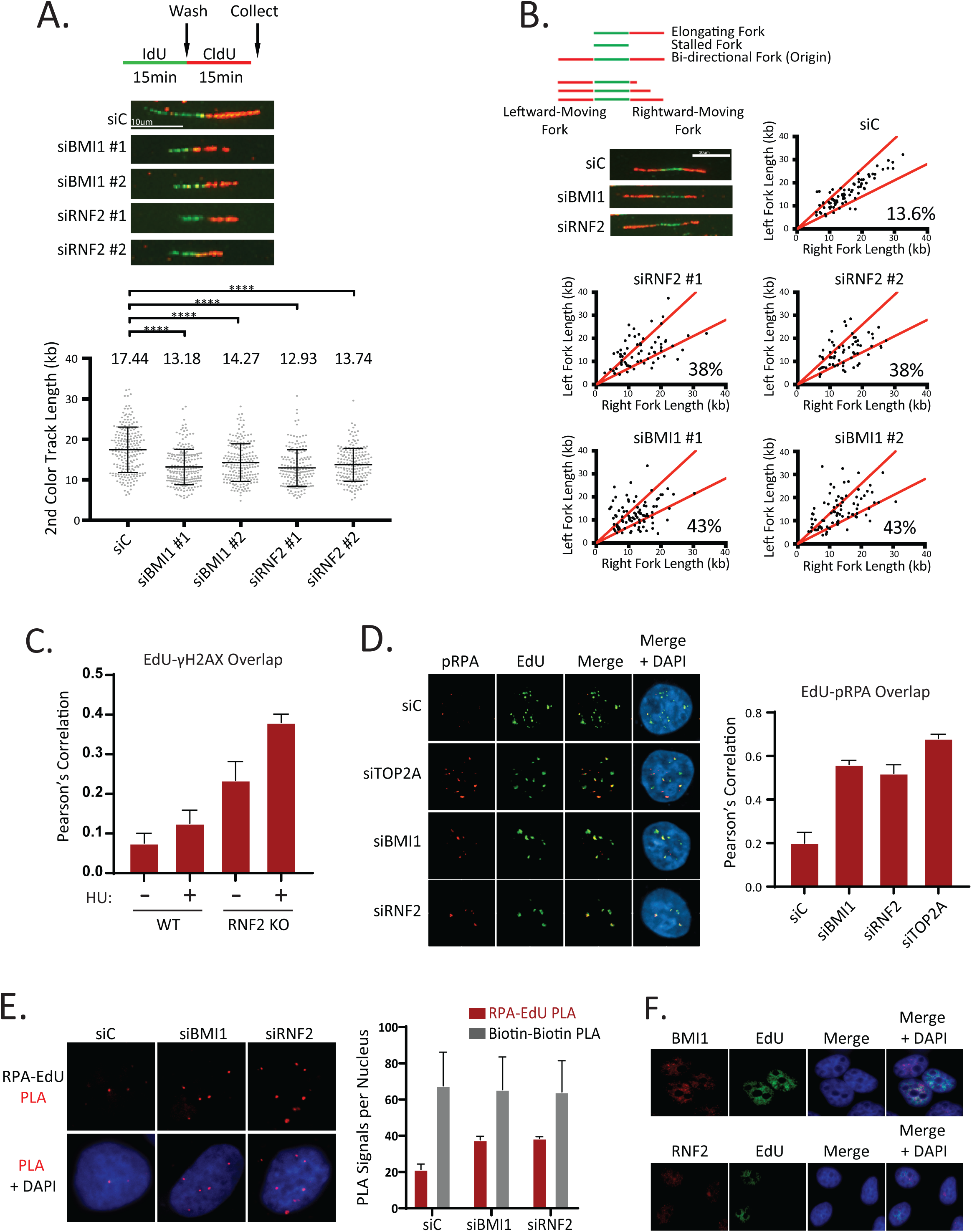
BMI1 and RNF2-deficiency causes increased replication fork stress. **A.** (Top) Schematic for measuring replication fork speed by the DNA fiber analysis. RPE1 cells were labeled sequentially with IdU and CldU for 15 minutes each. Representative DNA fibers in each siRNA-transfected sample. (Bottom) Quantification of 2^nd^ color track length in each siRNA-transfected sample. Numbers on top of the graph indicate average track length (N=180 from 3 biological replicates for each knockdown). **B.** (Top) Schematic for measuring bi-directional (or asymmetric) fork arrest in the DNA fiber assays. Representative images are shown. (Bottom) Knockdown of BMI1 or RNF2 increases replication fork asymmetry in RPE1 cells. Cells were labeled with IdU and CldU as in A. **C.** Co-staining of EdU and γH2AX showing increased intensities of γH2AX in EdU-positive cells when the RNF2 KO cells are treated with 2mM HU. The overlap between EdU and γH2AX is represented as the Pearson’s correlation coefficient. (N=50 from 2 biological replicates). **D.** (Left) Co-staining of EdU and p-RPA32 demonstrates that intermediate replication structures (e.g. ssDNAs) are increased in BMI1, RNF2 and TOP2A knock down T80 cells. (Right). Quantification of overlap between EdU and RPA32 by the Pearson’s correlation coefficient. (N=75 from 3 biological replicates). **E.** (Left) Representative images showing that the PLA signals between EdU and RPA32 are increased in U2OS cells depleted of BMI1 or RNF2. (Right) Quantification of average PLA signals per nucleus. Biotin-biotin PLA signals are unchanged under the given conditions. (N=50 from 3 biological replicates). **F.** Co-staining of BMI1 (Top) or RNF2 (Bottom) with EdU confirms these proteins are expressed in the nucleus during S phase.

Consistent with the observed replication stress, we found that BMI1 or RNF2 silencing led to increased γH2AX or RPA staining at EdU-labelled nascent replication forks (Figs 2C and 2D, respectively), which are indicatives of increased ssDNA formation and replication fork stalling. Knockdown of TOP2A also showed an increased association of RPA at forks, which served as a positive control. To further strengthen this finding, we employed a modified Proximity Ligation Assay (PLA)-based assay, which combines the Click chemistry-based labeling of nascent forks with EdU and measures the proteins association at nascent DNA (37). In this assay, U2OS cells were pulsed with 100uM EdU for 8 minutes before processing for the Click reaction and PLA (see methods for details). When BMI1 or RNF2 was knocked down, similar increases in the association of RPA at the nascent forks was observed (Fig 2E). Biotin-only controls showed equal staining, confirming equal reaction efficiencies and EdU labeling across the samples (S5 Fig). iPOND (isolation of proteins on nascent DNA) also showed increased RPA in RNF2-depleted cells (S6 Fig). BMI1 or RNF2 expression can be detected in EdU-labeled S phase cells (Fig 2F), supporting the S-phase dependent roles of these factors.

### Depletion of BMI1 or RNF2 causes transcription stress at CFSs and transcription-replication collisions

BMI1 has a role in repressing RNAPII elongation near damaged chromatin such as DSBs (31). We observed that RNF2 also has similar activities, which can be reversed by RNAPII elongation inhibitor DRB (S7 Fig). These results suggested that BMI1 and RNF2 suppress aberrant RNAPII activities near DNA breaks. Since CFSs are sites of DNA instability prone to breakages, we hypothesized that BMI1 and RNF2 also control RNAPII elongation at these sites. In the cells depleted of RNF2 by siRNA (Fig 3A) or CRISPR (Fig 3B), we observed increased RNAPII occupancy throughout the CFSs tested, as shown using anti-Rpb1 ChIP (P-Ser2; marker of elongating RNAPII). Endpoint PCR analysis showed consistent results (S8 and S9 Figs), and western blot confirms equal immunoprecipitation between the samples (S10 Fig). There was no detectable increase in the RNAPII occupancy in GAPDH, demonstrating the selective increases in CFSs. The increased RNAPII presence may indicate RNAPII experiencing increased arrest and pausing, or so-called transcription stress. The Flex1 region within FRA16D contains a high level of AT-rich sequence that can cause stalling of replication fork or transcription (5, 38), and it is possible that the transcriptional stress seen in RNF2-depleted cells may differ up or downstream of the Flex1 site. Unexpectedly, we observed increased RNAPII presence in all three sites tested in the RNF2 KO cells (Fig 3C; end-point PCR analysis showed consistent results in S11 Fig). Increased or aberrant transcription pausing or arrest are considered a source of transcription-replication collisions (TRCs), thus we wished to test incidences of TRC when BMI1 or RNF2 was depleted. Since the PLA can detect transient protein associations with high sensitivity, we explored the usage of PLA-based assays for detecting the physical association between the replisome and the largest subunit (Rpb1) of the RNAPII complex. We attempted to use the antibodies against MCM helicase subunits, but we could not detect any PLA signal when combined with several anti-RPB1 antibodies. However, we saw robust PLA signals when antibodies against PCNA and Rpb1 (p-Ser2) were used in BMI1 or RNF2 knockdown cells (Fig 3D). There were little signals in control siRNA-transfected cells, nor when either antibody was used alone. Interestingly, when RNAPII inhibitors DRB or alpha-amanitin were treated, the PLA signals were largely absent in the BMI1 or RNF2 knockdown condition (Fig 3E), suggesting that increased RNAPII elongation is responsible for the physical coupling between the two complexes. The PLA signals were also observed in T80 RNF KO cells, which were reversed by re-expressing RNF2 wild type (Fig 3F). We have extensively tested the authenticity of the PLA signals using other various gene knockdowns; knockdown of TCOF1 (nucleolar protein) or RNF20 (induces H2B ubiquitination) did not induce PLA signals. Knockdown of USP16, which deubiquitinates H2AK119-Ub (39), did not induce PLA signal either, possibly suggesting that the fine-tuning (ubiquitination and deubiquitination) of H2A-Ub may not be involved in the repression of TRC.

**Figure 3.**
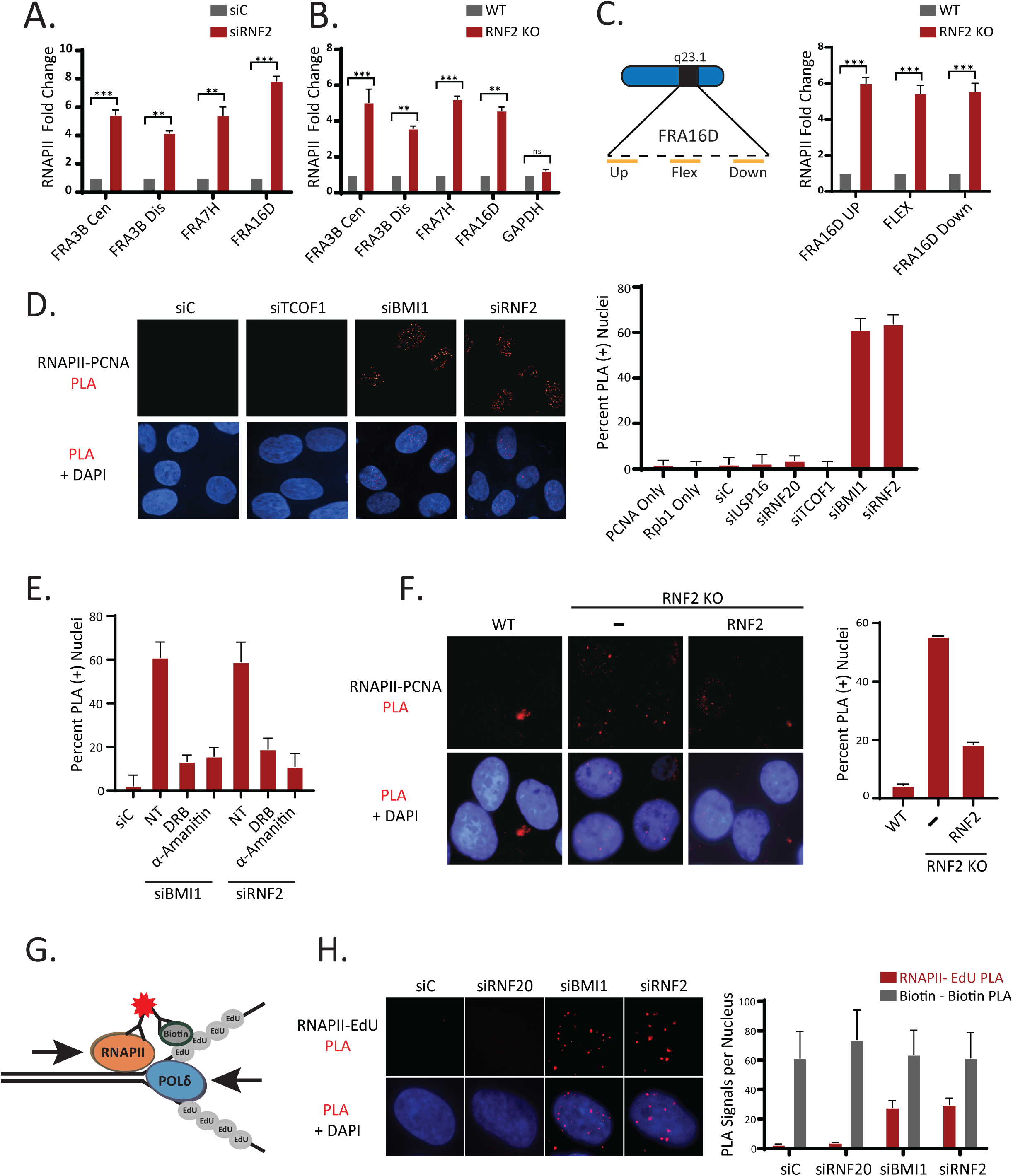
Irregular transcription is linked with transcription-replication conflicts in BMI1 and RNF2-deficient cells. **A-B.** ChIP using the Rpb1 (p-Ser2) antibody followed by qPCR amplification with the indicated primers demonstrates that the elongation of RNAPII is increased at the tested CFSs in RNF2 knockdown (siRNF2) (**A**) and KO (**B**) cells. (N=3 biological replicates; ***P <0.0005, **P <0.005). **C.** (Left) schematic of primer binding locations within the FRA16D locus (Right) ChIP using a Rpb1 (p-Ser2) antibody followed by qPCR amplification with indicated primers. (N=3 biological replicates; ***P < 0.0005, **P <0.005). **D.** (Left) Representative images demonstrating that the PLA signals between Rpb1 (p-Ser2) and PCNA is increased in T80 cells depleted of RNF2 or BMI1. (Right) Quantification of the percentage of PLA-positive nuclei under the indicated conditions (N=100 cells per condition from 3 biological replicates). **E.** The PLA signal between Rpb1 (p-Ser2) and PCNA is restored to normal levels by treatment with the transcriptional inhibitors DRB or α-Amanitin. Quantification of the percentage of PLA positive cells under the indicated conditions (N=100 nucleus per condition from 3 biological replicates). **F.** T80 RNF2 KO cells were transfected with 3xFLAG-RNF2 WT, fixed at 36 hours post-transfection, then analyzed for PLA as in D. The assays were done in triplicates (N=120 for each condition). **G.** Schematic presentation for detection of collisions between the replisome and RNAPII at nascent replication forks by PLA between biotin-labeled EdU and RNAPII. **H.** (Left) Representative images demonstrating that PLA signal between Rpb1 (p-Ser2) and EdU-labeled replication forks is increased in U2OS cells depleted of BMI1 and RNF2. (Right) Quantification of the average PLA signals per nucleus and biotin-only control PLA is shown. (N=50 from 3 biological replicates).

To further strengthen the finding of TRC, we used the modified PLA-based method using EdU labeling of nascent forks to detect the collisions (Fig 3G). The association between Rpb1 and EdU is absent in the control cells, but the signals were again significantly induced by knockdown of BMI1 or RNF2, but not by siRNAs targeting other genes (Fig 3H). Biotin-biotin antibody pair control showed equal staining (S12 Fig). Based on our extensive analysis, we conclude that the PLA assay faithfully represent the increase association (collision) of replisome and Pol II complex in BMI1 or RNF2-deficient cells.

### Fanconi Anemia proteins respond to R-loop-associated transcriptional stress in RNF2-deficient cells

As collisions of transcription and replication machineries are generally associated with R-loop accumulations (40), we tested whether RNF2 depletion causes an increased R-loop accumulation at the CFSs. R-loops can be detected by measuring the transient accumulation of the catalytically dead RNaseH1 (RNH1), a nuclease that detects and cleaves R-loops (41). Indeed, anti-V5-RNH1 ChIP assays showed that a catalytically inactive RNH1 (D210N) is more enriched at CFSs in the HU-treated cells compared to non-treated control cells (S13 Fig). Importantly, the mutant RNH1 is significantly more enriched at CFSs in RNF2 KO cells compared to control cells (Fig 4A), indicating that R-loops are indeed increased at CFSs in the absence of RNF2. Consistently, ChIP using anti DNA-RNA hybrid antibody (S9.6) showed similar results (Fig 4B). A series of reports have suggested that FA proteins engaged in resolving R-loops (see Discussion), and recent reports found that FANCD2 becomes enriched at CFSs when cells were challenged with replication stressors (18, 42). We thus tested if FANCD2 proteins are differentially enriched at CFSs in WT versus RNF2 KO cells. Anti-FANCD2 ChIP assays showed that FANCD2 is approximately 6 to 8 times more enriched at the tested CFSs in RNF2 KO compared to parental control cells (Fig 4C). To investigate in what extent the R-loops contribute to the FANCD2 accumulation at CFSs in RNF2 KO cells, we overexpressed RNH1 in the RNF2 KO cells and performed the anti-FANCD2 ChIP assays. We found that overexpressing RNH1 WT, but not RNH1 D210N, largely reduced the FANCD2 enrichment at CFSs in the RNF2 KO cells (Fig 4D), suggesting that the FANCD2 enrichment at CFSs in RNF2 KO cells are largely due to the increased R-loops. Further, we detected increased FANCD2 foci overlapping with RPA-coated single stranded DNA in RNF2 KO cells by immunofluorescence assay, which was partially reduced by overexpressing RNH1 (Fig 4E). To test if the increased R-loops are present at the replication forks and they are physically associated with the fork proteins, we immunoprecipitated the R-loops using the S9.6 antibody after crosslinking the cells with formaldehyde. We found that the eluate from RNF2 KO cells contained more fork-associated proteins FANCD2, FANCI, MCM7, and PCNA than the control cells (Fig 4F), supporting that the R-loops are increased at the forks in RNF2 KO cells.

**Figure 4.**
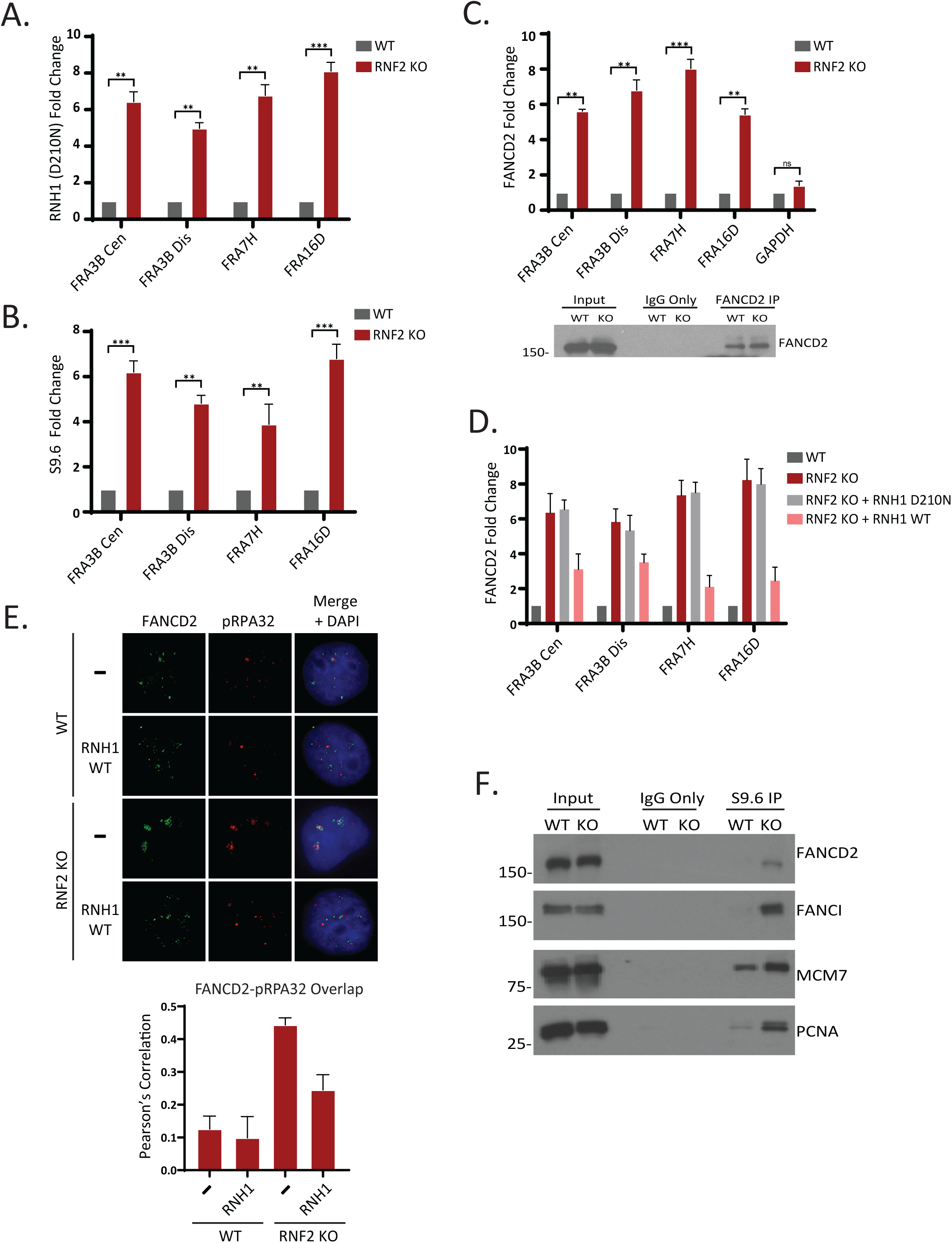
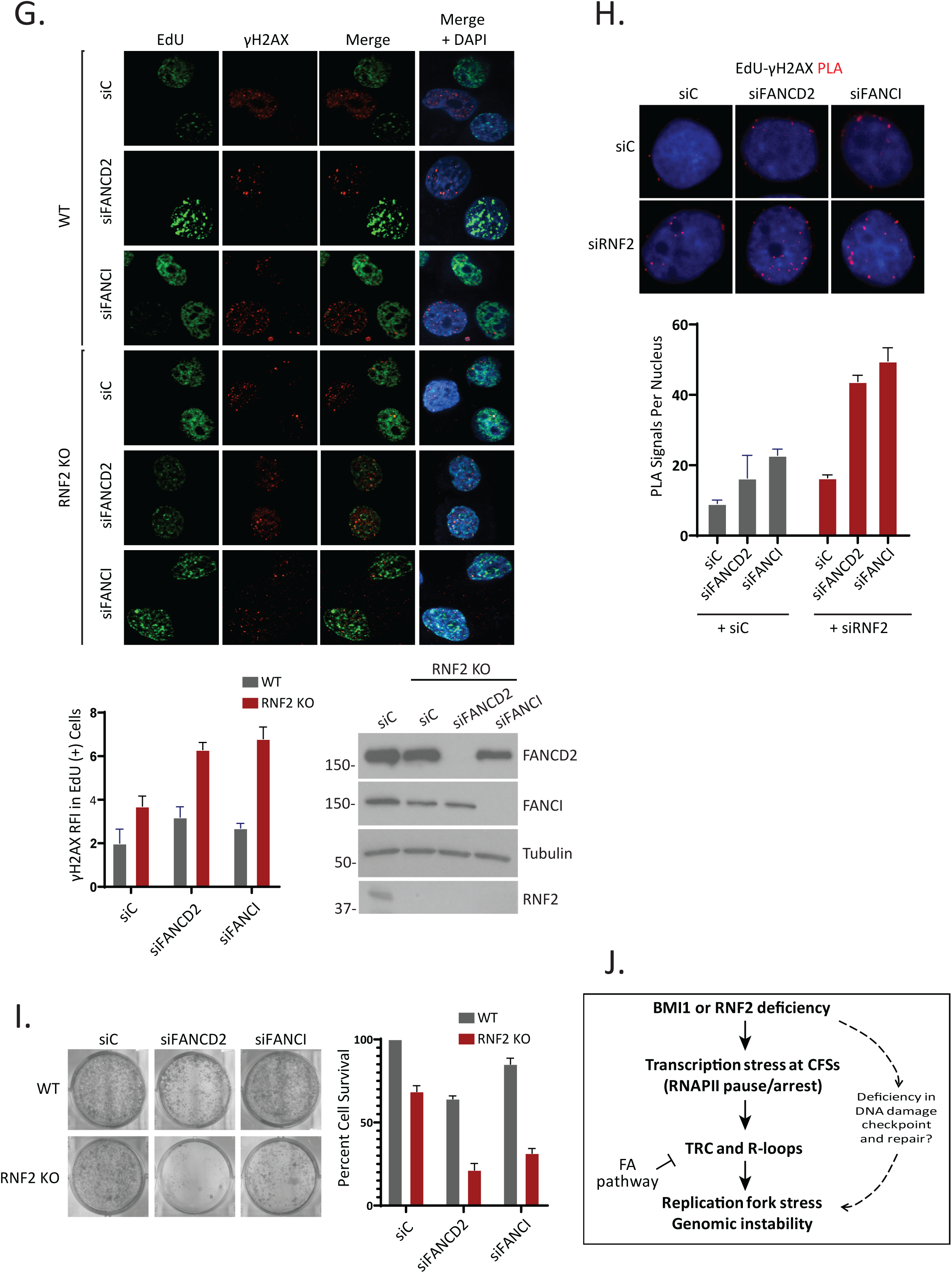
RNF2-deficient cells depend on the Fanconi Anemia fork-protective proteins for R-loop suppression, genome maintenance, and survival. **A.** T80 cells were transfected with pyCAG_RNaseH1_ D210N plasmid and subjected to ChIP with the anti-V5 antibody. qPCRs using indicated primers show that the R-loops are enriched at CFSs in the RNF2 KO cells (N= 3 biological replicates; ***P <0.0005, **P <0.005). **B.** ChIP using S9.6 antibody and amplification with the indicated primers by qPCR shows that R-loops are increased at CFSs in RNF2 KO T80 cells (N = 3 biological replicates, ***P <0.0005, **P <0.005). **C.** (Top) ChIP using FANCD2 antibody and amplification with the indicated primers by qPCR shows that FANCD2 is enriched at CFSs in RNF2 KO T80 Cells. (Bottom) Western blot confirming FANCD2 expression and IP efficiency in T80 WT and RNF2 KO cells (N=3 biological replicates; ***P <0.0005, **P <0.005). **D.** ChIP using FANCD2 antibody and amplification with the indicated primers by qPCR shows that FANCD2 enrichment at CFSs in RNF2 KO T80 cells is reduced by expressing exogenous RNH1 WT. There was no significant change upon expressing RNH1 D210N (N = 3 biological replicates). **E.** (Top) Representative images of FANCD2 and RPA foci in WT and RNF2 KO cells. Where indicated, cells were transfected with pyCAG_RNaseH1_ WT plasmid. (Bottom) Quantification of overlap between the FANCD2 and RPA signals by Pearson’s correlation (N=50 from 3 biological replicates). **F.** T80 WT and RNF2 KO cells were crosslinked, and the lysates were subjected to immunoprecipitation with the S9.6 antibody and the eluates were analyzed by western blots for indicated proteins. **G.** (Top) Representative images of EdU and γH2AX foci in WT and RNF2 KO T80 cells, where indicated FANCD2 and FANCI were also depleted by siRNA. (Bottom left) Quantification of the γH2AX RFI in EdU positive cells (N=75 from 3 biological replicates). (Bottom right) Verification of knockdown efficiency by western blot. **H.** (Top) Representative images showing the PLA signal between γH2AX and EdU-labeled replication forks is enhanced by the co-knockdown of RNF2 with either FANCD2 or FANCI in U2OS cells. siC indicates scrambled control siRNAs. (Bottom) Quantification of the number of PLA signals per nucleus under the indicated conditions (N= 50 from 3 biological replicates). **I.** Viability of the T80 RNF2 KO cells is decreased by the depletion of FANCD2 or FANCI by siRNA (N=6 biological replicates). **J.** Model for our findings.

Consistent with the perceived role of FANCD2 in relieving replication fork stress and mitigating R-loops, depleting FANCD2 with siRNA further increased the γH2AX foci in the RNF2 KO cells (Fig 4G). Depleting FANCI, a protein that forms a heterodimer with FANCD2, also led to similar increase in the γH2AX foci was observed in the KO cells, suggesting that FANCD2 and FANCI may act in a concerted manner. Similar results were obtained with FANCD2 and BMI1 knockdowns together (S14 Fig). The γH2AX-EdU PLA assay consistently showed similar increase when FANCD2 or FANCI are co-depleted with RNF2 (Fig 4H). These results suggest that FANCD2 and FANCI may act to suppress the R-loop-associated instabilities at CFSs in RNF2 or BMI1-depleted cells. Next, we investigated if the increased genomic aberrations correlated with cell death. siRNA-mediated depletion of either FANCD2 or FANCI further reduced the viability of the RNF2 KO T80 cells (Fig 4I). Altogether, these results suggest that there is an increase in R-loop formation in RNF2 KO cells, and the Fanconi Anemia proteins FANCD2 and FANCI are necessary to prevent the R-loop-associated genomic instability in RNF2 KO cells.

## Discussion

In this study, we provide evidence that Polycomb gene repressors BMI1 and RNF2 have a non-canonical role in suppressing transcription-replication collisions and R-loop suppression. We found that BMI1 or RNF2 deficiency causes increased replication stress, fork stalling, and CFS fragility. These phenotypes are associated with increased occupancy of RNAPII at CFSs, and physical collisions between RNAPII and replisome (or nascent forks). Consistent with the increasingly appreciated role of FANCD2 in R-loop binding and resolution, FANCD2 occupancy at CFSs is increased in RNF2 KO T80 cells, and is required to suppress genomic instability in RNF2 KO cells. Based on these data, we provide a model that BMI1 or RNF2 deficiency causes R-loop formation and TRCs at CFSs, which can be counteracted by the Fanconi Anemia proteins (Figure 4J).

The current notion is that transcription and replication occur in spatially and temporally separated domains (8). Whether this coordination is actively enforced by trans-acting factors, especially within CFSs that are prone to transcription stress, remains an important question. One known factor acting on the conflict resolution is RECQL5, a DNA helicase that associates with both RNAPII and PCNA (43–45); of note, RECQL5 depletion leads to uncontrolled elongation of RNAPII, with higher levels of RNAPII pausing and arrest globally at the transcription regions (45). We postulate that BMI1 and RNF2 may impose similar control over RNAPII elongation at CFSs. The increased occupation of RNAPII at CFSs (Figure 3) could be an indicative of increased transcriptional stress. At nuclease-induced DSBs, RNF2 depletion causes uncontrolled elongation of RNAPII that could be reversed by DRB (Figure S7). These results collectively suggested to us that RNAPIIs are “unleashed” in the absence of BMI1 or RNF2, causing transcription stress, and posing as hindrance to ongoing replisomes.

The genome maintenance role of BMI1 or RNF2 in replication-dependent context was also noted previously; BMI1 knockout MEFs cells show increased chromosome breakages when treated with replication stressors HU or APH (30), and RNF2 promotes replication elongation in pericentromeric region and S phase progression (46). Importantly, a recent work showed that RNF2 modulates the R-loop-associated transcription stress, in which overexpressing RNH1 could reverse the replication fork elongation deficiency in RNF2-depleted cells (47). Our findings add important new angles to this finding, by showing that RNF2 or BMI1 deficiency causes increased instabilities and transcription stresses at CFSs, and that the FA proteins act to mitigate the transcription stresses at CFSs. Thus, our work reveals the new cooperative relationship between the Polycomb proteins and the FA proteins in the CFS stability maintenance. RNF2 may act to repress the R-loop formation by inducing H2AK119-ub (47), which may create a state of chromatin where RNAPII progression is not permissive. Alternatively, as our previous work suggested that FACT-dependent RNAPII elongation is aberrantly regulated in BMI1-deficient cells (31), it is similarly possible that BMI1 and RNF2 controls the FACT-dependent RNAPII elongation at CFSs that could “smoothen” or buffer” the RNAPII-dependent transcription.

Our PLA analysis for TRC measurements suggest that physical association between RNAPII and replisome can occur. TRCs can occur in the form of head-on (HO)-oriented collisions or co-directional (CD)-oriented collisions. Studies found that genomic regions prone to the HO collisions are associated with R-loops (14, 48), and that the HO collisions causes activation of the ATR kinase (14). Based on the data that significant R-loop increase is seen in the RNF2-deficient cells, we extrapolate that the RNF2 deficiency may cause HO-oriented collisions. It is possible that under replication stress, BMI1 and RNF2 act to repress ongoing RNAPII elongation that could otherwise collide with newly initiated forks in HO-orientation. Further, treatment of an ATR inhibitor increased the cell death of BMI1 or RNF2 knockdown cells, as measured by sub-G1 apoptosing populations (S15 Fig). BMI1 depletion was also previously shown to increase the activation of the ATR-CHK1 kinases under replication stress condition (49), which might be relevant to our results.

Our work provides a functional link between the Polycomb proteins and the Fanconi Anemia DNA damage response pathway in suppressing genomic instability. Our data suggest that perturbed replication forks observed in BMI1 or RNF2-depleted cells may be at least partially salvaged by FANCD2. It is to be noted that BMI1 or RNF2 depletion still gives rise to replication stress in the presence of FA genes, suggesting that the stress burden may be too severe even with the normal FA gene functions. Our work also provides an example that the FA pathway can be activated by endogenously-triggered transcription stress (e.g. RNF2 mutation), in addition to commonly used drugs such as HU or APH. Apart from the well-established functions in resolving the DNA interstrand crosslinks, roles of the FA pathway preserving CFS stability and mitigating the R-loop-associated genome instability is growingly appreciated (16-20, 50, 51); in particular, FANCD2 localizes to sites of transcription (20), and purified FANCD2-FANCI heterodimer can directly bind to R-loops (17, 18). FANCD2 deficiency increases the replication stress at FRA16D region (19), and a genome-wide ChIP-sequencing analysis revealed the preferred accumulation of FANCD2 at CFSs under replication stress (18), supporting the importance of FANCD2 in CFS stability. A few studies may have suggested how FANCD2 facilitates the R-loop processing; FANCD2 may recruit RNA processing factors (50), or facilitate FANCM translocase activity (20). FANCD2 may also facilitate the recruitment of chromatin remodeler ATRX to CFSs (42) to resolve R-loop-mediated replication stresses (52). FANCD2 is necessary for recovery of perturbed replication forks through recruiting CtIP nuclease (53), which can facilitate the R-loop removal (54). FANCI is also known to activate dormant origin firing when the forks experience stress (55), therefore it is possible that the role of FANCI in the context of BMI1-RNF2 deficiency is to salvage the stalled forks by activating dormant origins within or nearby CFS. Future studies may provide further insights into the precise role of the FANCD2-FANCI heterodimer in resolving the R-loop-mediated replication stresses. Lastly, BMI1 knockout mice display significant defects in the hematopoietic system and bone marrow development (56, 57), a phenotype reminiscent of the Fanconi Anemia pathway deficiency. Although the phenotype could be majorly contributed by de-repression of BMI1 target genes (e.g. CDK inhibitors), we postulate that increased replication fork stress and CFS instability may also contribute.

Altogether, our work suggests that BMI1 and RNF2 bear a critical influence on the integrities of replication fork and CFSs, and emphasizes their tumor suppressive, rather than the often-perceived oncogenic, roles.

## Methods

### Cell lines, plasmids, and chemicals

T80 cells were grown in RPMI Medium supplemented with 10% FBS. HeLa, 293T, U2OS cells were grown in Dulbecco’s Modified Eagle’s Medium (DMEM) supplemented with 10% FBS. RPE1 cells were grown in DMEM F12 media supplemented with 10% FBS. RNF2 knockout T80 clones were generated using the CRISPR-Cas9 Double Nickase plasmid synthesized by Santa Cruz Biotechnology. U2OS cells stably expressing mCherry-LacI-Fok1 fusion protein that induces a DSB at a single genomic locus is previously described (58). pyCAG_RNaseH1_WT and D210N plasmids were a gift from Dr. Xiang-Dong Fu (Addgene plasmids #111906, # 111904). Hydroxyurea (AC151680050), Aphidicolin (61197-0010) and DRB (NC9855607) were purchased from Fisher Scientific. alpha-amanitin was purchased from Santa Cruz Biotechnology (sc-202440). IdU (I7125) and CldU (C6891) were purchased from Sigma Aldrich. ATR inhibitor (AZ20) was purchased from SelleckChem.

### RNAi

Cells were cultured in medium without antibiotics and transfected once with 20nM siRNA using the RNAiMAX (Invitrogen) reagent following the manufacturer’s protocol. The following siRNA sequences were synthesized by QIAGEN: RNF2 #1: AACGCCACUGUUGAUCACUUA, RNF2#3 UUGGGUUGCCACAUCAGUUUA, BMI1#1: AUGGGUCAUCAGCAACUUCUU, BMI1#2: CAAGACCAGACCACUACUGAA, FANCD2 GAGCCUGACAGAAGAUGCCUCCAAA, RNF20: ACGGGUGAAUUCCAAAGGUUA The following siRNA sequences were synthesized by Bioneer: FANCI: GACACCUUGUUAAAGGAC USP16: UGUGCAAGCUGUGCCUACA, TOP2A: GGUUGCCCAAUUAGCUGGA

### Western blots and antibodies

Cell extracts were run on an SDS-PAGE gel and then transferred to a PVDF membrane (Bio-Rad, Hercules, CA). Membranes were probed with primary antibodies overnight at 4°C. The membranes were then washed and incubated with either mouse or rabbit secondary antibody linked with horseradish peroxidase (Cell Signaling Technologies) and washed. The bound antibodies were viewed via Pierce ECL Western Blotting Substrate (Thermo Scientific). The following primary antibodies were used: α-BMI1, Ring1b (RNF2), MCM7, FANCI, RPA32, 53BP1 rabbit polyclonal antibodies are from Cell Signaling Technology. α-β-Tubulin and γH2AX mouse monoclonal antibodies are from Millipore. α-FANCD2, PCNA, and Cyclin A mouse monoclonal antibodies are from Santa Cruz Biotechnology. α-Biotin mouse monoclonal antibody and α-RPB1 (p-Ser2) rabbit polyclonal antibodies are from Abcam. α-V5 rabbit polyclonal antibody is from Invitrogen. α-DNA-RNA hybrid (S9.6) antibody is from Sigma Aldrich.

### Immunofluorescence

Cells were seeded in 12 well plates onto coverslips, indicated siRNA and damage treatments were applied. Media was removed from the wells, coverslips were washed twice with ice cold PBS and fixed for 10 minutes in the dark with cold 4% paraformaldehyde. The coverslips were washed twice with cold PBS and permeabilized for 5 minutes via incubation with 0.25% Triton and washed twice with cold PBS. Primary antibodies were diluted in PBS (1:300-1:500) and 30ul was applied to each coverslip before incubating for 1 hour in the dark, coverslips were washed twice with cold PBS. Secondary antibodies were diluted 1:1000 in PBS and 35ul was applied to each coverslip before incubating for 1 hour in the dark, coverslips were washed twice in PBS and placed onto glass slides. Vectashield mounting medium for fluorescence with DAPI (Vector Laboratories Inc) was used to stain nuclei. Images were collected by a Zeiss Axiovert 200 microscope equipped with a Perkin Elmer ERS spinning disk confocal imager and a 63x/1.45NA oil objective using Volocity software (Perkin Elmer). For detection of EdU-positive cells, cells were incubated with 10µM EdU for 15 minutes prior to fixing under normal growth conditions. After fixing and permeabilizing (as above), EdU was labeled with Alexa Fluor 488 Azide (Thermo Fisher) by a standard coper-catalyzed click reaction. Cells were co-stained for additional proteins where indicated, following our standard protocol (above). All fluorescence quantification was performed using ImageJ. To measure relative fluorescence intensity (RFI), single cells were manually selected, the integrated density was measured and corrected to account for background in the image. The density measurements were normalized with a value of 10 corresponding to the brightest reading. Pearson’s overlap correlations were obtained with the use of the “Colocalization finder” plugin for ImageJ. Full color images were imported into ImageJ and the channels were split into blue, red, and green; the red and green channels were analyzed and the degree of colocalization was determined.

### Proximity Ligation Assay (PLA)

Proximity ligation assays were preformed using the Duolink kit from Sigma Aldrich; cells were grown in a 12 well format on coverslips. Cells were fixed and permeabilized according to the standard immunofluorescence protocol (previously described), primary antibodies were added at a 1:500 dilution in PBS and incubated for 1 hour at room temperature. PLA minus and plus probes were diluted 1:5 in the provided dilution buffer, 30ul of the probe reaction was added to each coverslip and incubated for 1hr at 37C; the coverslips were washed twice with buffer A. The provided ligation buffer was diluted 1:5 in water, the ligase was added at a 1:30 dilution; the ligation reaction was left at 37°C for 30 minutes before washing twice with wash buffer A. The provided amplification buffer was diluted 1:5 in water before adding the provided polymerase at a 1:80 ratio, the amplification reaction was left at 37°C for 100 minutes, the reaction was quenched by washing twice with buffer B. The coverslips were mounted on slides with DAPI containing mounting medium. For EdU-PLA, cells were seeded glass coverslip and treated with indicated siRNA for 72 hours. Cells were pulsed with 100uM EdU for 8 minutes before fixing with 4% paraformaldehyde for 10 minutes, washed with PBS twice and permeabilized with 0.25% Triton X-100 for 5 minutes. Cells were then washed twice with PBS. Click reaction buffer (2mM copper sulfate, 10uM biotin azide, 100mM sodium ascorbate) was prepared fresh and added to the slides for 1 hour at room temperature. Cells were washed three times with PBS. Primary antibodies were diluted in PBS and added to slides for 1 hour at room temperature. Cells were washed with PBS twice. PLA reaction was then carried out as described above.

### Chromatin Immunoprecipitation

Cells were crosslinked with 1.42% formaldehyde for 10 minutes in the dark at room temperature (RT), the crosslinking was quenched by adding 125mM Glycine for 5 minutes in the dark at RT. Crosslinked cells were washed and harvested by scraping. Cells were lysed for 10 minutes on ice with the FA lysis buffer (50mM HEPES, 140mM NaCl, 1mM EDTA, 1% Triton X-100, 0.1% Sodium Deoxycholate). Lysates were sonicated at 45% amplitude 8 times for 10 seconds each, with 1 minute rest on ice between pulses. Inputs are collected and the lysates were incubated with the indicated antibodies at a 1:200 concentration overnight at 4°C. Protein G agarose beads were added to the lysates for ∼3 hrs, beads were washed 3 times with the FA lysis buffer prior to elution. To elute DNA, 400ul of Elution buffer (1% SDS, 100mM Sodium Bicarbonate) was added to the beads, then rotated at RT for 2 hours. The eluate was collected and incubated with RNase A (50ug/ml) for 1 hr at 65°C, followed by proteinase K (250ug/ml) overnight at 65°C. The DNA is purified with the PCR purification Kit (Bioneer) following the manufacture’s instructions. For the detection of R-loops, wild type and RNF2 KO T80 cells were transiently transfected with pyCAG_RNaseH1_ D210N plasmids, then cell pellets were harvested in ∼36 hours, lysed and sonicated as described above. The lysates were subjected to IP with the anti-V5 antibody (1:200 dilution) and processed as described above. For the crosslink-IP for western blot (Figure 4F), the procedure was the same except that the proteins were eluted by boiling in the SDS buffer.

### qPCR

All qPCR experiments were performed on an appliedbiosystems QuantStudio3 thermocycler using amfiSure qGreen Q-PCR master mix (GenDEPOT Q5603-001). All qPCR reactions were 50ul in volume and contained 15ng of template DNA. PCR cycles consisted of 35 cycles of 95°C denaturation for 15s followed by annealing/extension for 1 minute at 60°C, measurements were acquired after each cycle. Fold change quantification was preformed using the ΔCT of the untreated sample and the experimental sample for each primer set assuming the product was doubled for each cycle. The specify of amplification was confirmed by running products on agarose gels as well as melt curve analysis following every qPCR cycle. The Cq confidence of all samples quantified was greater than 0.98. Primers used for CFS amplification were: Primers used for CFS amplification were: FRA3B Central FW: 5’-tgttggaatgttaactctatcccat -3’, FRA3B Central RV 5’-atatctcatcaagaccgctgca -3’ FRA3B Distal FW: 5’-caatggcttaagcagacatggt -3’, FRA3B Distal RW: 5’-agtgaatggcatggctggaatg -3’, FRA7H FW: 5’-taatgcgtccccttgtgact -3’, FRA7H RV: 5’-ggcagattttagtccctcagc -3’, FRA16D (UP) FW: 5’-tcctgtggaagggatattt -3’, FRA16D (UP) RV: 5’-cccctcatattctgcttcta -3’, FRA16D (FLEX) FW: 5’ – gatctgccttcaaagactac – 3’,FRA16D (FLEX) RV: 5’ – caaccaccattctcactctc – 3’, FRA16D (DOWN) FW: 5’ – cagattcctttttctcattg – 3’,FRA16D (DOWN) RV: 5’ – gttaggtctacattttcagt – 3’, GAPDH FW-5’ – ccctctggtggtggcccctt – 3’GAPDH RV-5’ – ggcgcccagacacccaatcc – 3’

### DNA Fiber Analysis

DNA fibers were prepared as described previously (59). Briefly, cells were pulsed with 50µM IdU and 100µM CldU for times indicated in each experiment. After trypsinization, cells were washed and resuspended at 1 × 10^6 /mL in cold PBS, 2uL were plated onto a glass slide, and lysed with 10µL lysis buffer (0.5% SDS, 200mM Tris-HCl pH 7.4, 50mM EDTA) for 6 min. Slides were tilted at a 15 degree angle to allow DNA spreading, and then fixed for 3 min in chilled 3:1 methanol:acetic acid. The DNA was denatured with 2.5 N HCl for 30 min, washed in PBS, blocked for 1hr in 5% BSA in PBS with 0.1% Triton X-100. Slides were stained for 1 hour with primary antibodies, washed 3X in PBS, stained for 30 min with secondary antibodies, washed 3X in PBS and dried. Coverslips were mounted with Prolong antifade reagent and sealed. Slides were imaged with Keyence BZ-X710 microscope. Image analysis was done with ImageJ. A minimum of 60 fiber lengths were measured for each independent experiment measuring track length, and analysis shows the pool of three independent experiments (biological replicates). Track lengths were calculated by converting µm measured in ImageJ to kb using the conversion 1µm = 2.59 kb. For asymmetric fork analysis, the left and right fork lengths were measured from bidirectional origin events, and asymmetry calculated as number of origins with lengths greater than +/− 30% variation from equal length over the total number of events. Anti-BrdU antibody (ab6326) (CldU) was purchased from Abcam. Anti-BrdU antibody (347580) (IdU) was purchased from BD Biosciences. Goat anti-Rat IgG (H+L) Cross-Adsorbed Secondary Antibody, Alexa Fluor 594 (A-11007) and Goat anti-Mouse IgG (H+L) Cross-Adsorbed Secondary Antibody, Alexa Fluor 488 (A-11001) were purchased from Thermo Fisher.

### Clonogenic survival assay

Cells were seeded into 24 well plates (∼10 cells per visual field) and treated with indicated siRNA for 48hours. UV irradiation (254nm) was applied using the Stratalinker UV crosslinker, 2400, then the cells were allowed to grow for 10∼14 days. The cells were fixed with a 10% methanol, 10% Acetic acid solution for 15minutes at room temperature, followed by staining with crystal violet. Sorensen buffer (0.1M sodium citrate, 50% ethanol) was used to extract the stain, then the colorimetric intensity of each solution was quantified using Gen5 software on a Synergy 2 (BioTek, Winooksi, VT) plate reader (OD at 595nm). Error bars are representative of 3 independent experiments.

### Cell Cycle Analysis

U2OS or T80 cells were transfected with siRNAs for 72 hours, followed by treatment with HU or ATRi where indicated. Cells were harvested and fixed with 70% Ethanol for 1 hour in darkness, washed with PBS and incubated with Propidium Iodide (50ug/ml), RNase (25ug/ml) and Triton X-100 (1%) for 1 hour. Cell cycle analysis was carried out in Accuri C6 Flow Cytometer and data was analyzed using BD Accuri C6 Software.

## Acknowledgement

We thank Dr. Catherine Freudenreich for helpful discussions and suggestions. We thank Drs. Meera Nanjundan for providing T80 cells, and Roger Greenberg for the PTuner263 cell line. pyCAG_RNaseH1_WT and D210N plasmids were gifts from Dr. Xiang-Dong Fu through Addgene. We thank Robert Hill and Charles Szekeres for technical support for confocal microscope and Flow cytometry, respectively.

---

**Figure S1.**
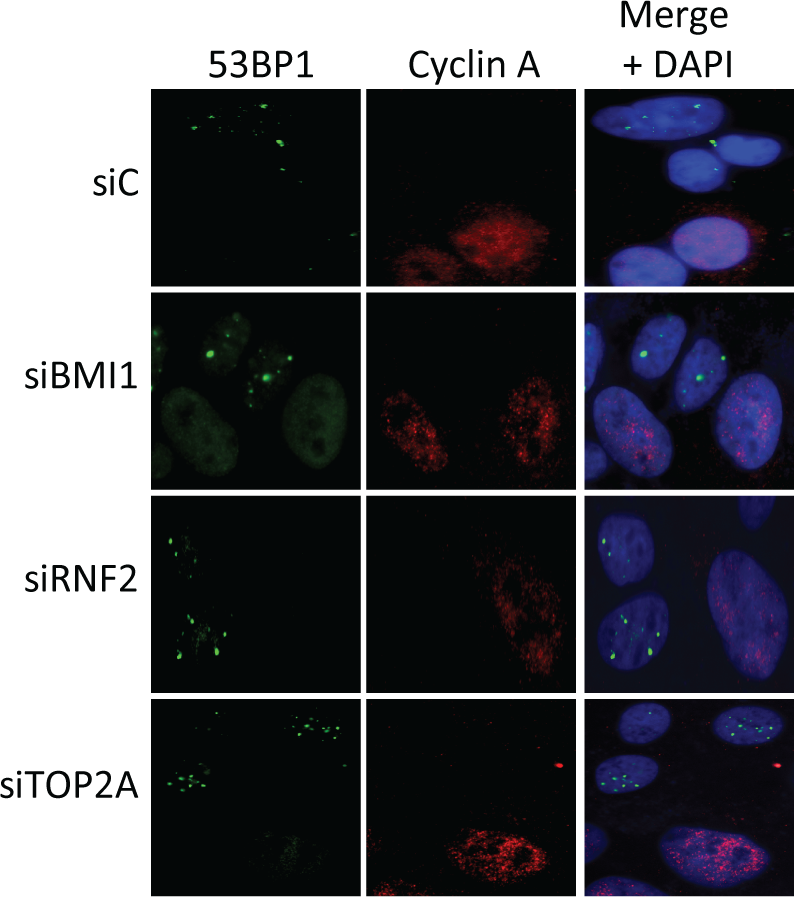
Representative images of U2OS cells stained with 53BP1 and Cyclin A following transfection with indicated siRNAs followed by treatment of 2mM HU for 16 hours. (Quantification is shown in Figure 1A).

**Figure S2.**
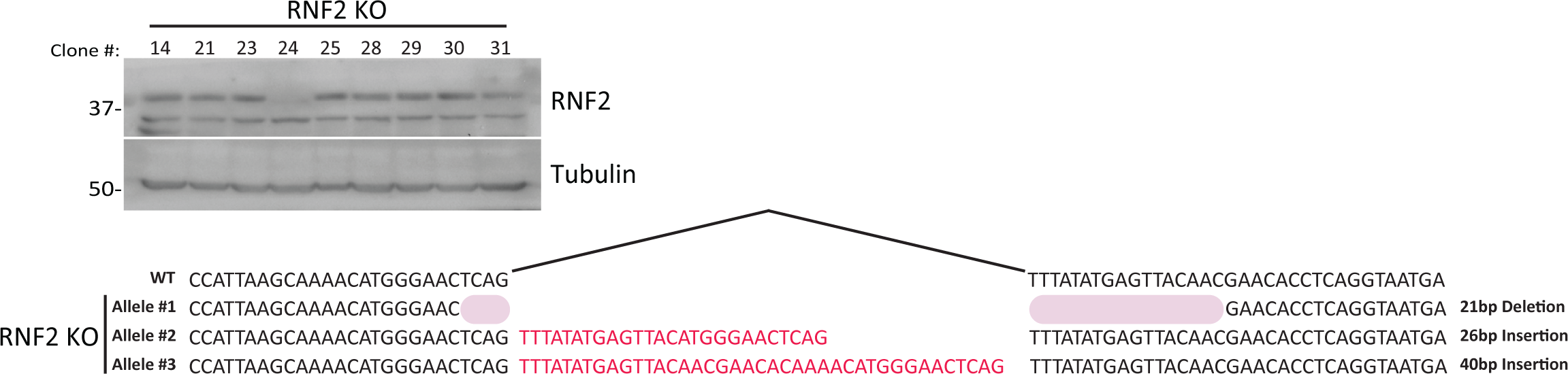
(Top) Western blot screening identifies the clone #24 as true T80 RNF2 CRISPR KO clone. (Bottom) Sequencing analysis of the RNF2 KO clone #24. Of the three alleles for RNF2 in T80, two contain frame shift insertions and one contains an in frame deletion.

**Figure S3.**
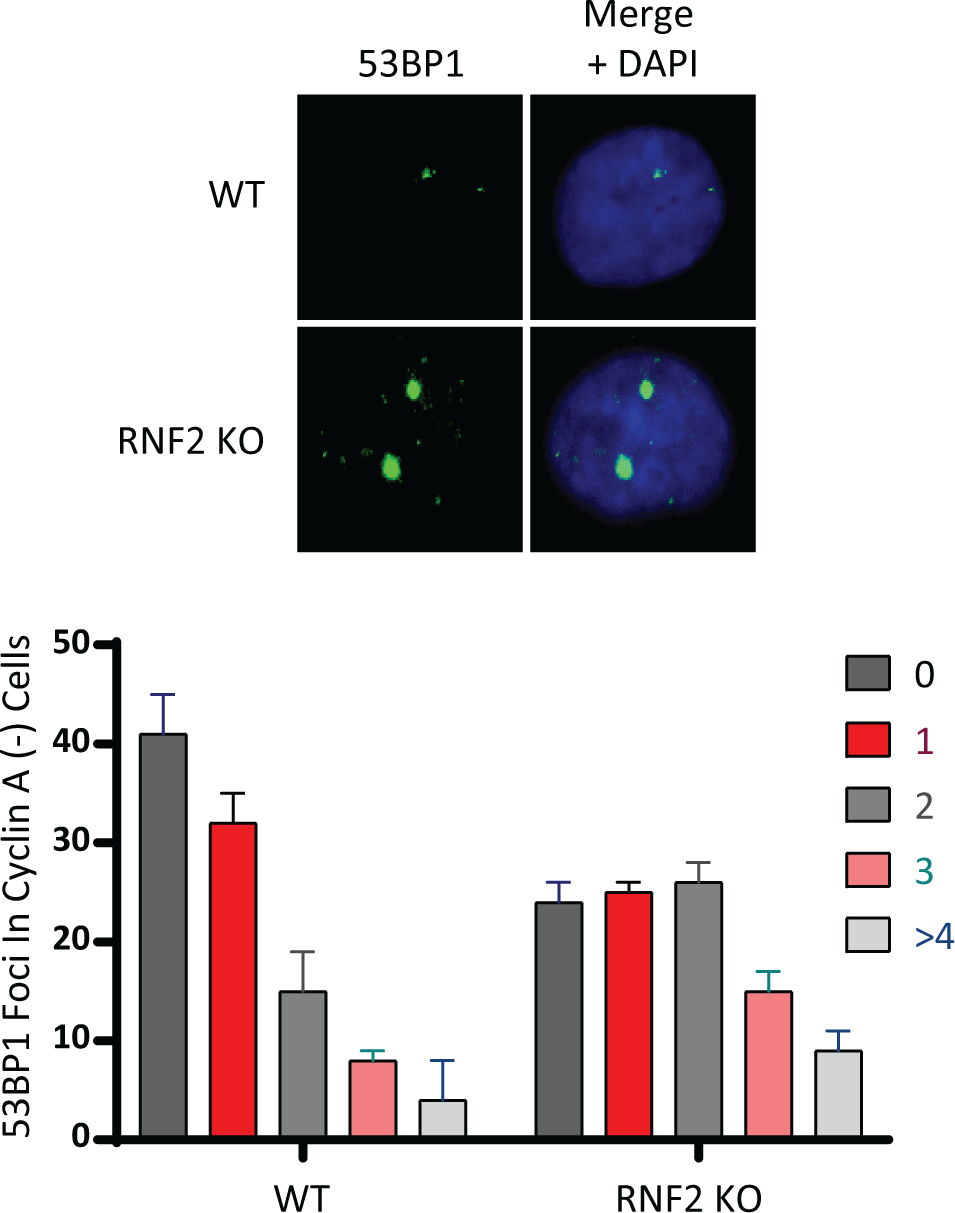
(Top) Representative images of T80 wild type and RNF2 KO cells stained with 53BP1. (Bottom) Quantification of the number of 53BP1 bodies per nucleus under the indicated conditions. (N=50 from 3 biological replicates)

**Figure S4.**
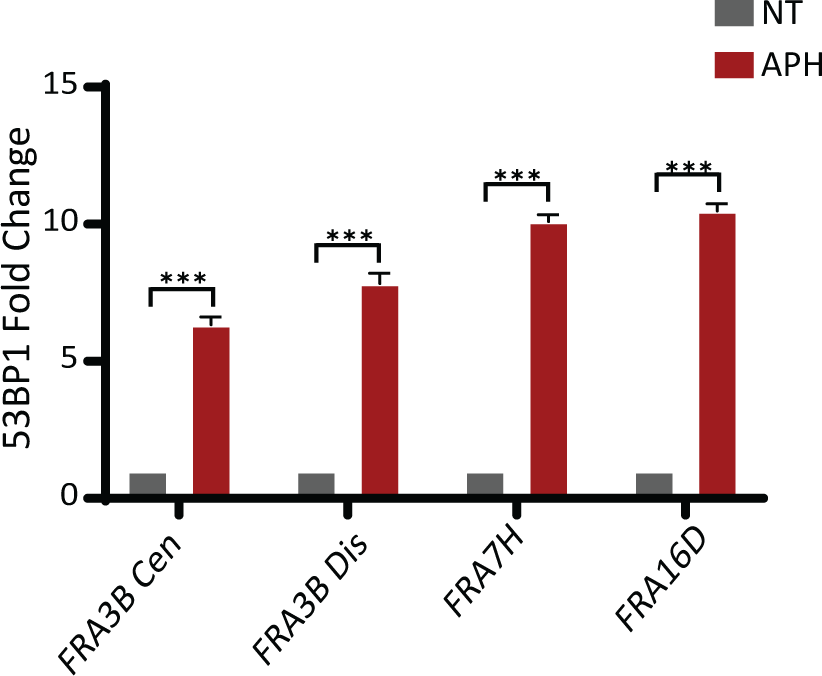
qPCR quantification of anti-53BP1 ChIP in wild type T80 cells with or without treatment with 0.4µM Aphidicolin for 16 hours. (N=3 biological replicates; ***P < .0005,**P <.005, *P <.01).

**Figure S5.**
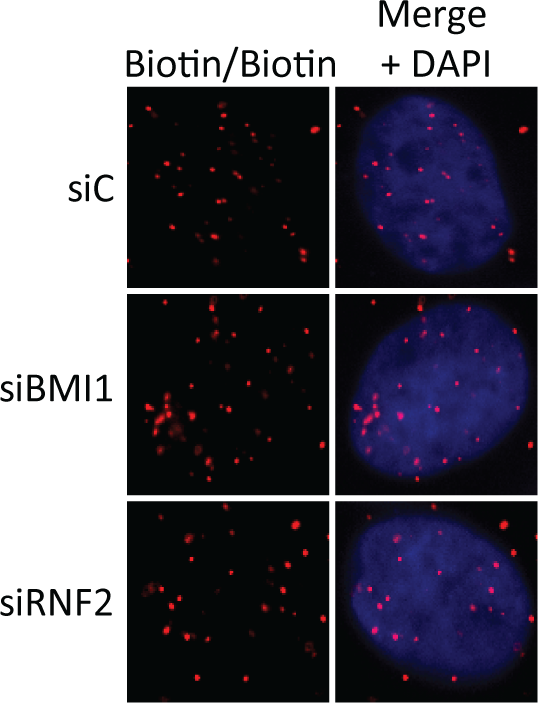
U2OS cells were labeled with EdU and subjected to a Click reaction with azide biotin. The cells were probed with mouse and rabbit biotin antibodies and used for a PLA reaction to determine if the extent of EdU labeling was equal among the conditions. (N=3 biological replicates). These cells were set up simultaneously with sample probed for pRPA32 and EdU (Figure 2E).

**Figure S6.**
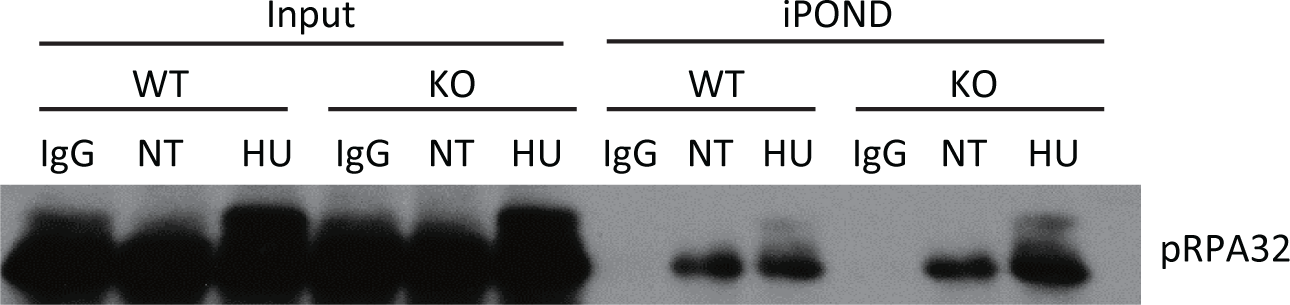
The isolation of protein on nascent DNA (iPOND) assay demonstrates that phosphorylated RPA is enriched at the replication fork in T80 RNF2 KO cells. Where indicated cells were treated with 2mM HU for 16 hrs.

**Figure S7.**
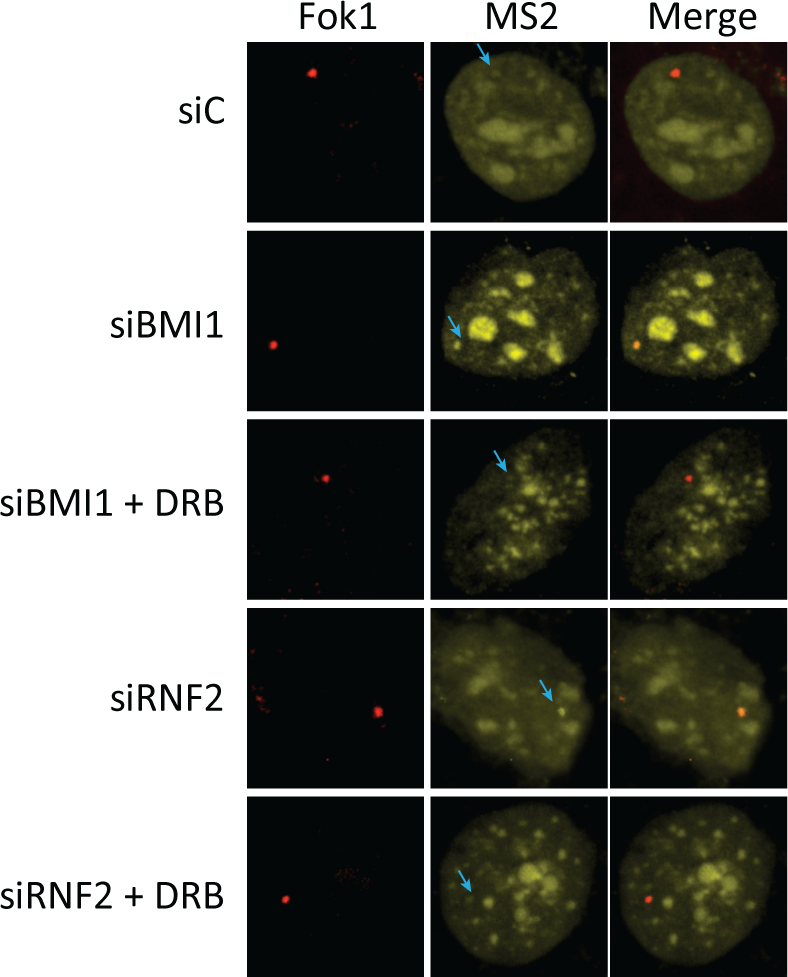
Assays using the pTuner263 transcriptional reporter cell line demonstrates that transcriptional output (measured by YFP-MS2 signal) at double strand breaks (marked by the FOK1 endonuclease) is unregulated in RNF2 and BMI1 knockdown cells. This effect is reversed by treatment with 50uM DRB.

**Figure S8.**
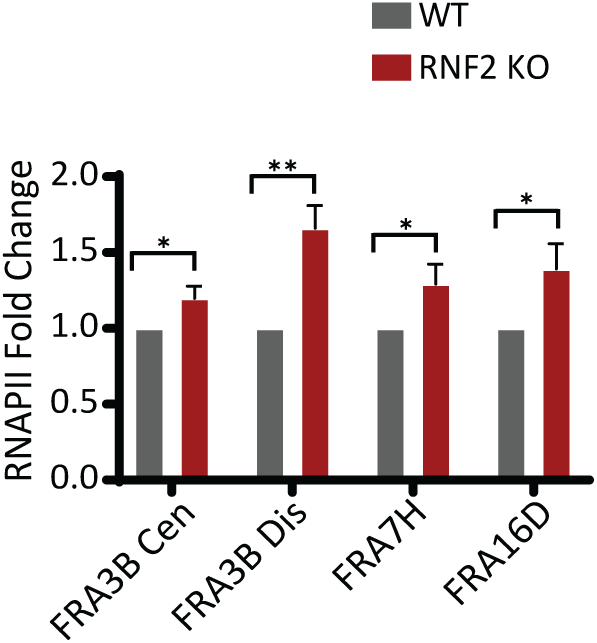
Quantification of end-point band intensity of RNAPII ChIP. T80 wild type and RNF2 KO cells were IP’ed with anti-Rpb1 (P-Ser2) antibody and the bound DNA was amplified with the indicated primers. (N=3 biological replicates).

**Figure S9.**
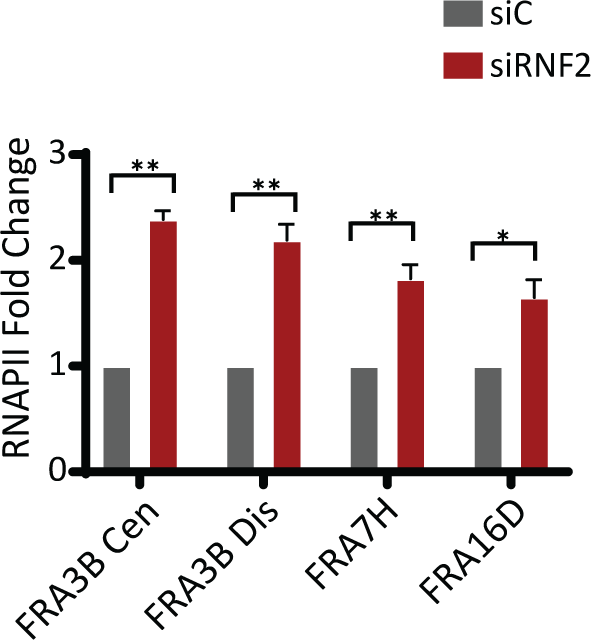
Quantification of end-point band intensity of RNAPII ChIP. siControl and RNF2-knockdown T80 Cells were IP’ed with anti-Rpb1 (P-Ser2) antibody and the bound DNA was amplified with the indicated primers (N=3 biological replicates; **P <.005, *P <.01).

**Figure S10.**
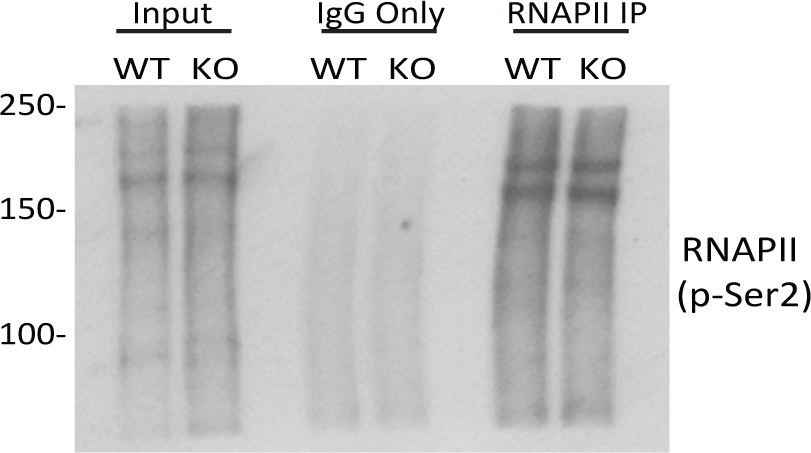
Western blot confirming that expression and IP of Rpb1 under the ChIP conditions was equal between the T80 WT and RNF2 KO cells.

**Figure S11.**
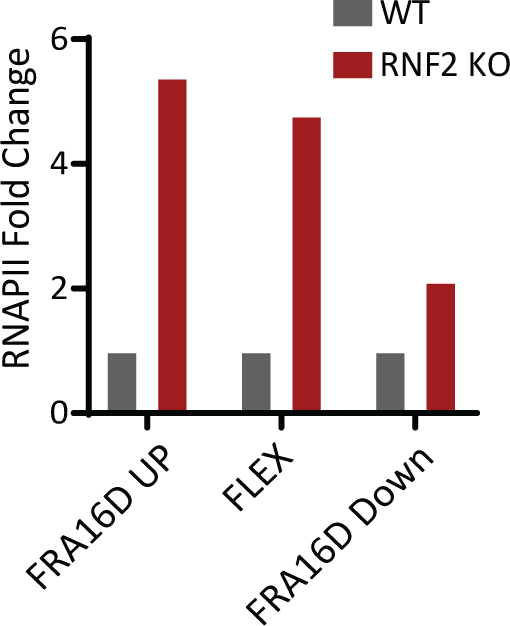
Quantifications for the end-point band intensity of RNAPII ChIP from T80 wild type and RNF2 KO cells. (Amplification with 3 primer sets within the FRA16D region.)

**Figure S12.**
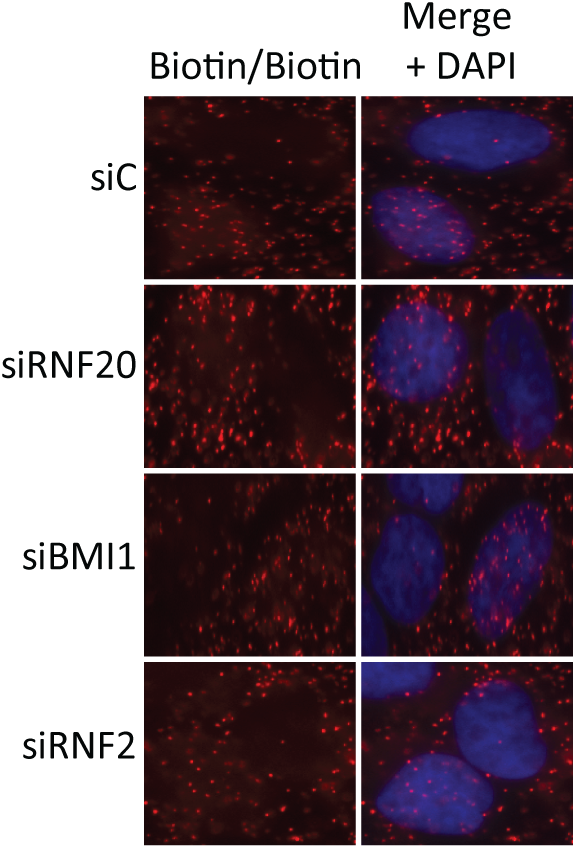
U2OS cells were labeled with EdU and subjected to the Click reaction with azide biotin. The cells were probed with mouse and rabbit anti-biotin antibodies and used for PLA reactions to determine if the extent of EdU labeling was equal between all conditions. (N=3 biological replicates). These cells were set up simultaneously with sample probed for Rpb1 and EdU (Figure 3H).

**Figure S13.**
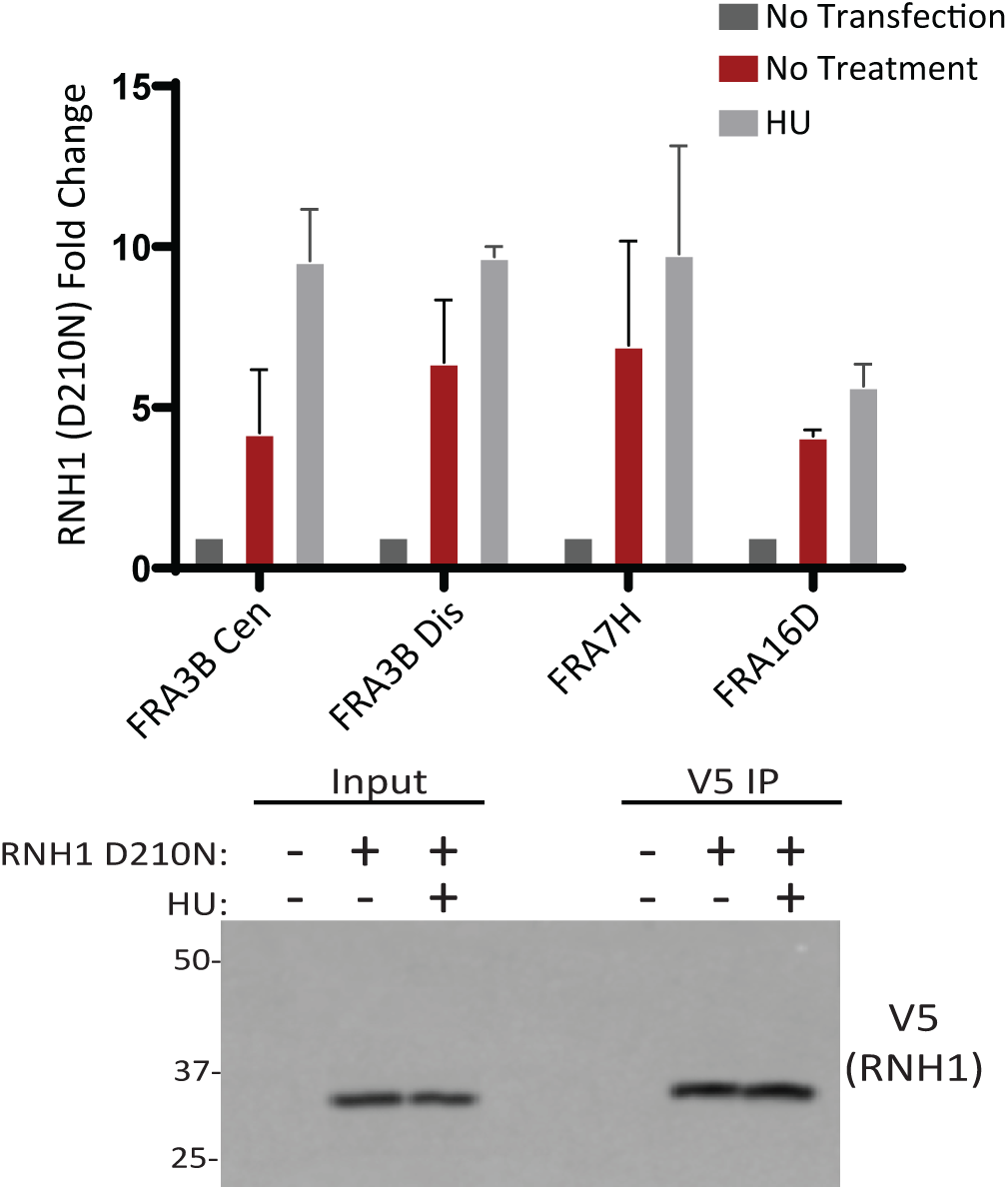
(Top) Quantification of end-point ChIP assay from T80 cells transfected with pyCAG_RNaseH1_ D210N. Cells were subsequently treated with HU (2mM) and IP’ed with anti-V5 antibody (N= 2 biological replicates). (Bottom) Anti-V5 western blot confirming the RNH1 expression and IP efficiency.

**Figure S14.**
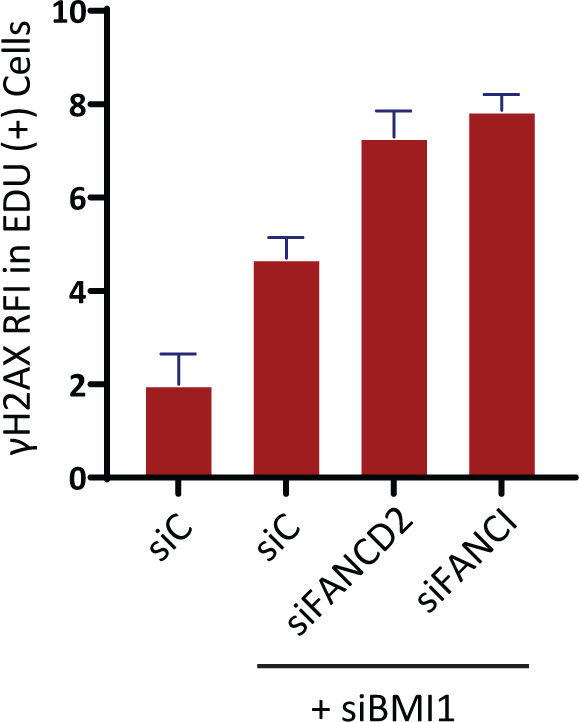
Quantification of γH2AX RFI in T80 cells depleted of BMI1 by siRNA. Where indicated, FANCD2 and FANCI were co-depleted by siRNAs. (N=50 from 3 biological replicates)

**Figure S15.**
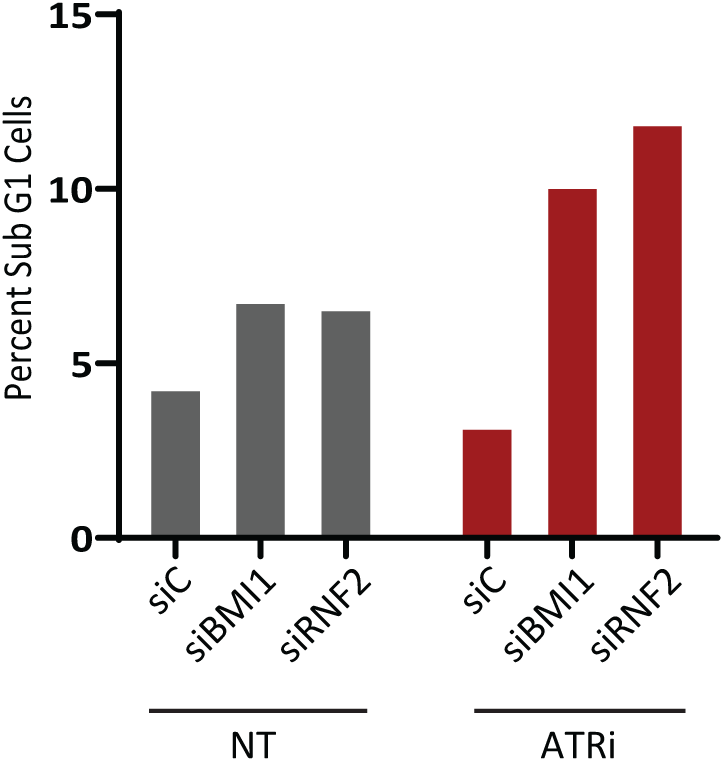
DNA content analysis by Flow cytometer shows that the percentage of sub-G1 cells is increased when BMI1 or RNF2 knockdown cells are co-treated with an ATR inhibitor (AZ20; 100nM, 16 hour treatment).

